# A potent and selective inhibitor for the modulation of MAGL activity in the neurovasculature

**DOI:** 10.1101/2022.05.04.490688

**Authors:** Alicia Kemble, Benoit Hornsperger, Iris Ruf, Hans Richter, Jörg Benz, Bernd Kuhn, Dominik Heer, Matthias Wittwer, Britta Engelhardt, Uwe Grether, Ludovic Collin

## Abstract

Chronic inflammation and blood–brain barrier dysfunction are key pathological hallmarks of neurological disorders such as multiple sclerosis, Alzheimer’s disease and Parkinson’s disease. Major drivers of these pathologies include pro-inflammatory stimuli such as prostaglandins, which are produced in the central nervous system by the oxidation of arachidonic acid in a reaction catalyzed by the cyclooxygenases COX1 and COX2. Monoacylglycerol lipase hydrolyzes the endocannabinoid signaling lipid 2-arachidonyl glycerol, enhancing local pools of arachidonic acid in the brain and leading to cyclooxygenase-mediated prostaglandin production and neuroinflammation. Monoacylglycerol lipase inhibitors were recently shown to act as effective anti-inflammatory modulators, increasing 2-arachidonyl glycerol levels while reducing levels of arachidonic acid and prostaglandins, including PGE_2_ and PGD_2_. In this study, we characterized a novel, highly selective, potent and reversible monoacylglycerol lipase inhibitor (MAGLi 432) in a mouse model of lipopolysaccharide-induced blood–brain barrier permeability and in both human and mouse cells of the neurovascular unit: brain microvascular endothelial cells, pericytes and astrocytes. We confirmed the expression of monoacylglycerol lipase in specific neurovascular unit cells *in vitro*, with pericytes showing the highest expression level and activity. However, MAGLi 432 did not ameliorate lipopolysaccharide-induced blood–brain barrier permeability *in vivo* or reduce the production of pro-inflammatory cytokines in the brain. Our data confirm monoacylglycerol lipase expression in mouse and human cells of the neurovascular unit and provide the basis for further cell-specific analysis of MAGLi 432 in the context of blood–brain barrier dysfunction caused by inflammatory insults.

## INTRODUCTION

The endocannabinoid system encompasses a vast network of G-protein-coupled receptors, endogenous signaling ligands and rate-limiting serine hydrolases responsible for endocannabinoid synthesis and degradation [1]. Cannabinoid receptors 1 and 2 (CB1 and CB2) are the most abundant and well-characterized cannabinoid receptors [2], although the endocannabinoid system also features several orphan receptors. Anandamide (AEA) and 2-arachidonyl glycerol (2-AG) are the two major endocannabinoid signaling lipids in the central nervous system (CNS). They both act as full agonists for CB1, whereas only 2-AG also acts as a full agonist for CB2. Both agonists are produced in the CNS, but 2-AG levels are ^~^170 fold higher than AEA levels in the murine brain, so 2-AG is responsible for most endocannabinoid signaling in that organ [3].

Monoacylglycerol lipase (MAGL) is the key rate-limiting enzyme that hydrolyzes 2-AG in the brain. It is a cytosolic membrane-bound serine hydrolase with two known isoforms of ^~^33 and ^~^35 kDa in mice and humans. MAGL activity is responsible for approximately 85% of 2-AG hydrolysis in the mouse CNS [4], the remainder hydrolyzed by α,β-hydrolase domain-containing protein 12 (ABHD12), ABHD6 and fatty acid amide hydrolase (FAAH). The latter is also primarily responsible for the hydrolysis of AEA. The direct byproducts of 2-AG hydrolysis by MAGL include large pools of glycerol and most of the arachidonic acid produced in the brain, which is then available for conversion to prostaglandin by the cyclooxygenases COX1 and COX2. In contrast, phospholipase A_2_ (PLA_2_) is the major enzyme responsible for arachidonic acid production in peripheral tissues, through the hydrolysis of membrane-bound phospholipids [5,6]. The genetic deletion or acute pharmacological blockade of MAGL in wild-type mice increases the levels of 2-AG and achieves a rapid and sustained reduction in the levels of arachidonic acid. Furthermore, the levels of 2-AG in the brain increased by ^~^10-fold in *MAGL^-/-^* mice, reflecting a 90% decrease in the rate of 2-AG degradation [7]. MAGL inhibitors therefore provide an effective anti-inflammatory approach targeting the arachidonic acid/prostaglandin axis in the CNS.

The provision of 2-AG confers neuroprotective effects by reducing blood–brain barrier (BBB) permeability, hippocampal cell death and inflammatory cytokine release in mouse models of traumatic brain injury and closed head injury [8,9]. Inhibiting MAGL activity in the brain is also beneficial in the aforementioned mouse models of brain injury and stroke by significantly elevating 2-AG levels while reducing the levels of arachidonic acid and its downstream metabolites, including prostaglandins and leukotrienes [8–12]. Furthermore, the pharmacologic inhibition of MAGL ameliorates lipopolysaccharide (LPS)-induced inflammation by reducing the production of pro-inflammatory cytokines and the associated vascular permeability [13]. We therefore investigated the role of MAGL and its inhibition at the cellular level of the BBB using human *in vitro* and mouse *in vivo* models.

The BBB is a dynamic vascular complex composed of brain microvascular endothelial cells (BMECs), which tightly regulate paracellular transport though tight and adherens junctions. These cells regulate the ability of solutes in the blood to enter the CNS at the capillary level. The BBB is part of a tightly coordinated cellular network known as the neurovascular unit (NVU) in which endothelial cells are ensheathed by pericytes and embedded in a shared endothelial basement membrane and further surrounded by astrocytic end feet processes and a second parenchymal basement membrane which together form the glia limitans. Pericytes are specialized mural cells that interact directly with the microvascular endothelial cells while the glia limitans provides an additional layer of coverage. Both pericytes and astroytes provide support by maintaining junctional proteins and regulating capillary vasodilation, blood flow and immune cell extravasation [14,15]. NVU dysfunction is an early pathological hallmark associated with multiple sclerosis, Alzheimer’s disease and Parkinson’s disease, in which microvascular dysfunction can promote immune cell infiltration, oxidative stress, impaired transport and clearance of inflammatory proteins, demyelination and/or neuronal loss [16–21]. The alteration of endocannabinoid signaling has also been reported in many of these neurological disorders, supporting the further investigation of endocannabinoid-mediated target therapies.

Although MAGL activity has been studied extensively in neurons and glia, little is known about MAGL expression and activity specifically at the level of the microvasculature and its role in the regulation of BBB permeability [12,22,23]. An astrocyte-specific *MAGL* knockout mouse model showed that MAGL is the enzyme primarily responsible for 2-AG hydrolysis in these cells, generating large pools of arachidonic acid that lead to prostaglandin production in the CNS [7]. The analysis of mouse cerebellar brain slices *ex vivo* suggests that PGE_2_ secreted by astrocytes regulates pericyte contraction and, in turn, the vasodilation and constriction of associated microvessels [24,25]. However, the inhibition of MAGL in NVU cells (BMECs, pericytes and astrocytes) is incompletely understood.

Enzyme inhibitors targeting the endocannabinoid system (including inhibitors of FAAH and MAGL) have been developed as therapeutics to attenuate prostaglandin-induced inflammation. Several MAGL inhibitors have been described [11,26,27], but most act irreversibly by forming covalent bonds with the catalytic Ser122 residue of MAGL[28]. The chronic administration of irreversible MAGL inhibitors in rodents can lead to CB1-mediated side effects, reflecting prolonged exposure to high levels of 2-AG and the subsequent desensitization of CB1 receptors, hindering clinical development [29]. This effect was also observed in *MAGL^-/-^* mice [29–31]. Furthermore, irreversible hydrolase inhibitors often lack selectivity, leading to potential adverse effects that can be fatal in some cases [32]. Therefore, the development of highly selective reversible MAGL inhibitors could improve the pharmacological control of enzyme activity and reduce off-target and adverse effects.

Here, we describe the characterization of a non-covalent, potent and selective MAGL inhibitor (MAGLi 432) in human and mouse brain lysates. MAGLi 432 (CAS no. 2361575-20-2)[33] is a bicyclopiperazine derivative optimized from a screening hit, which achieves high exposure in the mouse brain following intraperitoneal (i.p.) administration. We investigated the species-dependent expression profile of *MAGL* mRNA and MAGL protein in cultured microvascular endothelial cells, pericytes and astrocytes, and observed the modulation of cell-specific activity in human cells. We also used a mouse model of LPS-induced BBB disruption to investigate whether the administration of MAGLi 432 can ameliorate BBB permeability to 70-kDa dextran tracers and fibrinogen, and its ability to reduce the level of pro-inflammatory cytokines.

## MATERIALS AND METHODS

### Structure determination and refinement of human MAGL complexed with MAGLi 432

Human MAGL (residues 1–303) with mutations Lys36Ala, Leu169Ser and Leu176Ser [34] was purchased from Cepter Biopartners (Nutley, NJ, USA). Before crystallization, the protein was concentrated to 10.8 mg/mL. Crystallization trials based on sitting drop vapor diffusion were carried out at 21 °C. Crystals appeared within 2 days in solvent (0.1 M MES pH 6.5, 10% PEG MME5K, 12% isopropanol). Crystals were soaked for 16 hrs in the crystallization solution supplemented with 10 mM MAGLi 432.

For data collection, crystals were flash cooled in liquid nitrogen with 20% ethylene glycol added to the soaking solution as a cryo-protectant. X-ray diffraction data were collected at a wavelength of 0.9999 Å using an Eiger2X 16M detector at beamline X10SA of the Swiss Light Source (Villigen, Switzerland). Data were processed using XDS and scaled with SADABS (Bruker, Billerica, MA, USA). The crystals belong to space group C222_1_ with cell axes of a = 91.88 Å, b = 127.43 Å and c = 60.46 Å and diffract to a resolution of 1.16 Å. The structure was determined by molecular replacement with PHASER [35] using the coordinates of PDB entry 3PE6- as a search model. Fo-Fc difference electron density map was used to place MAGLi 432. The structure was refined using the CCP4 suite [36] and BUSTER [37], and model building done with COOT [38]. Data collection and refinement statistics are summarized in **Table S1**. The coordinates of the complex structure have been deposited in the Protein Data Bank under accession number PBD7ZPG.

### Primary cell cultures

Primary mouse brain microvascular endothelial cells (pMBMECs), pericytes and astrocytes originated from CD-1 mice. The pMBMECs (CD-1023) and astrocytes (CD-1285) were obtained from CellBiologics (Chicago, IL, USA). The pMBMECs were maintained in complete mouse endothelial cell medium from the CellBiologics W Kit (M1168), whereas astrocytes were maintained in phenol-free astrocyte medium – animal (1831-prf) from ScienCell (Carlsbad, CA, USA). Mouse brain vascular pericytes (ScienCell; M1200) were maintained in phenol-free pericyte medium – mouse (ScienCell; 1231-prf). All cells were maintained up to passage 4.

Primary human brain cortical pericytes (ScienCell; 1200) were maintained in phenol-free pericytes medium (ScienCell; 1201-prf), whereas primary human brain cortical astrocytes (ScienCell; 1800) were maintained in phenol-free astrocyte medium (ScienCell, 1801-prf). Brain microvascular endothelial cells, hCMEC/D3 cells [39] (Merck, Darmstadt, Germany; SCC066) were cultured in EGM-2 endothelial cell growth medium-2 from the BulletKit (Lonza, Basel, Switzerland; CC-3162). Pericytes and astrocytes were used for experiments up to passage 4, whereas hCMEC/D3 cells were used up to passage 40.

### Animals

Male CD-1 mice, 8–9 weeks of age, were acquired from Charles River (Lyon, France). All mice were housed individually in ventilated cages in a temperature-controlled environment with a 12-h photoperiod and were allowed food and water *ad libitum*. All procedures were conducted under the approval of the Veterinary Office of the Canton of Basel, Switzerland (License 2901).

### LPS-induced BBB disruption

MAGLi 432 was dissolved in the vehicle solution (a 1:1:8 mixture of DMSO, polysorbate 80 and 0.9% (w/v) NaCl). LPS from *Escherichia coli* O111:B4 (Sigma-Aldrich, St Louis, MO, USA; L2630) was dissolved in 0.9% NaCl. Mice were randomized to one of three treatment groups (1 = NaCl + vehicle, 2 = LPS + MAGLi 432, 3 = LPS + vehicle) and were injected i.p. with either 0.9% NaCl or 1 mg/kg LPS, followed 30 min later by the vehicle solution or 2 mg/kg MAGLi 432. Injections were performed for three consecutive days. On the third day, mice were euthanized 4 h after treatment by the i.p. administration of 150 mg/kg pentobarbital in 0.9% NaCl. Mice were then perfused with ^~^20 mL heparin (5 U/mL in phosphate-buffered saline (PBS), pH 7.4). Brains (divided sagittally into two hemispheres) and plasma were collected and immediately frozen at –80 °C.

### In vivo permeability assays and immunofluorescence

Mice were injected in the tail vein with 200 μL fluorescein isothiocyanate (FITC)-labeled dextran (Thermo Fisher Scientific, Waltham, MA, USA; D1820) prepared as a 25 mg/mL solution in 0.9% NaCl. The dye was allowed to circulate for 15 min before the mice were euthanized and perfused with ^~^10 mL heparin (5 U/mL in PBS, pH 7.4) then 10 mL 2% paraformaldehyde (PFA) in PBS (pH 7.4). The brains were fixed in 2% PFA at 4 °C overnight and then stored in PBS at 4 °C.

Whole brains were embedded in 2% agarose blocks and 100-μm sagittal sections were prepared using a vibratome (Zeiss, Oberkochen, Germany). In all subsequent steps the samples were protected from light. Sections were then incubated for 1 h with BlockAid Blocking Solution (Thermo Fisher Scientific; B10710) at room temperature, followed by incubation with a primary antibody specific for CD31 (BD Pharmingen, Franklin Lakes, NJ, USA; 550274) overnight at 4 °C. Goat anti-rat IgG (H+L) Alexa Fluor 555 (Invitrogen, Thermo Fisher Scientific; A-21434) was used as the secondary antibody, followed by nuclear staining with DAPI. Sections were imaged with a Pannoramic p250 Slide Scanner (3DHistech, Budapest, Hungary) and analyzed using CaseViewer v2.64.

### Measurement of brain and plasma fibrinogen

One cerebral hemisphere (without the cerebellum or olfactory bulb) was homogenized in radioimmunoprecipitation (RIPA) assay buffer (Sigma-Aldrich; R0278) containing Pierce protease and phosphatase inhibitors (Thermo Fisher Scientific; 88669). The tissue in RIPA buffer was placed in 7-mL Precellys homogenization vials (Bertin Instruments, Montigny-le-Bretonneux, France; P000940-LYSK0), prefilled with ceramic homogenization beads and was subjected to three 10-s pulses at 6000 rpm in a Precellys Evolution homogenizer (Bertin Instruments). The lysates were centrifuged twice at 16,300 x g for 15 min and the supernatants were retained for analysis. For plasma sample collection, blood was collected by cardiac puncture and placed in EDTA-coated tubes, which were centrifuged at 3220 x g for 5 min. The plasma was collected and centrifuged again at 17,530 x g for 5 min to remove any residual red blood cells. Brain homogenates (1:30 dilution) and plasma samples (1:100,000 dilution) were analyzed for fibrinogen levels using an Aviva Systems Biology (San Diego, CA, USA) ELISA kit (GWBBB0BA2).

### RNA extraction, cDNA generation and droplet digital qRT-PCR

RNA was extracted from sections of frozen mouse brain cerebrum representing each treatment group (n = 6 per group) using the Maxwell 16 LEV simplyRNA Tissue Kit (Promega, Ipswich, MA, USA; AS1270). RNA was extracted from cultured cells using the Maxwell 16 Cell LEV Total RNA Purification Kit (Promega; AS1225). Corresponding cDNA was prepared using the SuperScript III First-Strand Synthesis System (Invitrogen).

Commercially available TaqMan assays (**Table S2**) for droplet digital PCR were prepared using the One-Step RT-ddPCR Advanced Kit for Probes (Bio-Rad, Hercules, CA, USA; 1864022). Each 20-μL reaction mix consisted of 10 μL ddPCR Supermix for probes (no dUTP) 1 μL of each target probe, 1 μL of reference probe and up to 8 μL of cDNA or water, making a final volume of 13 μL. The reaction mix was then partitioned into droplets using a QX100 Droplet Generator (Bio-Rad). Target and reference probe fluorescence amplitudes for each droplet were analyzed using a QX200 Droplet Reader (Bio-Rad) for digital absolute quantification. Each sample was analyzed in duplicate.

### In vitro investigation of MAGLi 432 and LPS

Human astrocytes, pericytes and hCMEC/D3 cells were grown to confluence in T25 flasks. MAGLi 432 was prepared as a 10 mM stock solution in DMSO. LPS was solubilized in HBSS and was prepared as a 1 mg/mL stock solution. One day before the assay, cells were cultured in serum-free media. Cells were exposed to 1 μM MAGLi 432, 0.01% DMSO, no treatment (serum-free medium only), 0.01% HBSS, 50 ng LPS or 100 ng LPS for 6 h. Media and cell pellets were immediately collected and stored at −80° for LC-MS analysis or cells were lysed in activity based protein profiling (ABPP) lysis buffer (20 μL/mL 1M HEPES, 2 μL/mL 1M DTT, 1 μL/mL 1M MgCl, 0.5 μL/mL <u>(25U/ul)</u> Benzonase) for ABPP and western blot analysis.

### Competitive ABPP for MAGL selectivity

Homogenates of whole mouse brains or human cortical brain tissue (BioIVT, Westbury, NY, USA) without disease pathology were prepared in ABPP lysis buffer. Tissue in ABPP lysis buffer was homogenized as described above, and the lysate supernatants were retained. We incubated 50 μg of the lysates with 0.5 μL DMSO, 10 μM MAGLi 432 or 10 μM JZL 184 for 25 min, and then with 0.5 μL of 200 nM TAMRA-FP (green) or 5 μM MAGL-specific fluorescent probe [40] for 30 min, protected from light. The reaction was stopped by adding 12.5 μL deionized water, 12.5 μL 4x sample buffer (Thermo Fisher Scientific; NP0007) and 5 μL reducing agent (Thermo Fisher Scientific; NP0004). We loaded 25 μL of cell lysate solution into the wells of NuPAGE 10% Bis-Tris gels (Invitrogen; NP0301BOX) and run for 55 min at 200 V. We measured in-gel fluorescence on the ChemiDoc MP System (Bio-Rad) using Cy3 and Cy5 filters. The membrane was then stained with Coomassie Brilliant Blue for total protein quantification (**Fig. S1**).

### Competitive ABPP/western blot for MAGL dose response

Brain lysates (50 μg) as described above were incubated for 25 min with 0.5 μL DMSO or ascending doses of MAGLi 432 (1 nM, 10 nM, 100 nM, 1 μM or 10 μM). Lysates were then incubated for a further 30 min with 0.5 μL of the 5 μM MAGL-specific fluorescent probe [40]. The reaction was stopped and electrophoresis was carried out as described above. Proteins were transferred to nitrocellulose membranes (Invitrogen; IB23001) using the iBlot 2 System (Invitrogen). Membranes were blocked in 5% skimmed milk in PBS (pH 7.4) containing 0.01% Tween-20 (Sigma-Aldrich, P1379) for 1 h at room temperature and were incubated overnight at 4 °C with primary rabbit polyclonal antibodies against mouse MAGL (Abcam, Cambridge, UK; ab180016) or human MAGL (LSBio, Seattle, WA, USA; LS-C482667-200). On the following day, membranes were washed and incubated with the goat anti-rabbit IgG HRP-labeled secondary antibody (PerkinElmer, Waltham, MA, USA; NEF812001EA) for 1 h at room temperature. Band intensity was measured using the ChemiDoc MP System (Bio-Rad).

### ABPP/western blot for MAGL activity in neurovascular cells

Mouse (pMBMECs, primary astrocytes and pericytes) and human (hCMEC/D3 cells, primary astrocytes and pericytes) NVU cells were grown to confluence before dissociation with TrypLE and centrifugation. Cell pellets were resuspended in ABPP lysis buffer on ice for 30 min, and were vortexed until lysis was complete. Cell lysates (20 μg) were incubated for 25 mins with 0.5 μL of the 5 μM MAGL-specific fluorescent probe, and 10 μg of each lysate was loaded onto a NuPAGE 10% Bis-Tris gel for in-gel fluorescence and western blot analysis as described above.

### ABPP/western blot for MAGL activity in mouse cortical lysates

Brain homogenates were prepared from one cerebral hemisphere as described above and the lysis supernatant was collected for ABPP and western blot analysis as described above. In each case, the total protein content of the sample was measured using a BCA assay. MAGL activity probe bands and MAGL protein intensity bands were quantified using ImageLab (Bio-Rad). TAMRA-FP selectivity studies were normalized to Coomassie Brilliant Blue band intensity. All other western blots were normalized to β-actin protein band intensity.

### LC-MS assessment of mouse brain lysates

Deep frozen mouse brains were placed into 7 ml Precellys vials prefilled with ceramic homogenization beads (Bertin Instruments, P000940-LYSK0) and homogenized in methanol using a Precellys Evolution Homogenizer (Bertin Instruments) at 3 x 10 s, 6000rpm giving a final concentration of 100 mg brain/ml methanol. Samples were cleaned up by protein precipitation and aliquots were diluted 1:2 with methanol containing internal standards. Undiluted precipitates were used to measure AEA, PGE_2_ and PGD_2_ levels, whereas precipitates were diluted 1:100 to measure 2-AG and arachidonic acid (in each case 2 μL of each sample was injected). Calibration samples were prepared by spiking the pooled brain precipitate with 10-fold concentrated calibration standards serially diluted in methanol. The internal standards for arachidonic acid, 2-AG, AEA, PGE_2_ and PGD_2_ were AA-d8, 2-AG-d5, AEA-d8, PGE_2_-d4 or PGD_2_-d9, respectively (all from Cayman Chemicals, Ann Arbor, MI, USA). Calibration was achieved by linear regression with weighting 1/y and excluding zero. Absolute sample concentrations were calculated by dividing the peak area ratio analyte/internal standard by the slope of the calibration curve.

### LC-MS assessment for primary cell lysates

Cell pellets and supernatants stored at –80 °C were resuspended in 50 μL distilled water, briefly vortexed and sonicated for 15 min, before adding 100 μL methanol and repeating the vortex and sonication steps. The samples were then centrifuged at 18,000 × g for 10 min to precipitate the protein. The clean precipitate was diluted 1:2 with methanol containing internal standards and 2 μL of each sample was directly injected for analysis. Cell supernatants were diluted 1:2 with methanol containing internal standards and vortexed briefly before 2 μL of each sample was directly injected for analysis. Calibration samples were prepared for every tissue by spiking cell precipitates or supernatants with 5-fold concentrated calibration standards in methanol.

### In situ hybridization

Control human brain cortical tissue in paraffin blocks was sourced from an in-house tissue repository. Mouse brain cortical tissue was obtained from CD-1 mice not exposed to any treatment. Brains were removed from freshly euthanized mice and placed in 10% formalin at 4 °C for 24 h before embedding in paraffin blocks and preparing 4-μm tissue sections using a microtome (Leica Microsystems, Wetzlar, Germany). We detected *MAGL* mRNA *in situ* using the RNAscope 2.5 Assay for Ventana Discovery Ultra system (ACD Bio, Minneapolis, MN, USA) and the RNAscope VS Universal HRP Reagent Kit (Red) (ACD Bio; 323250). Mounted tissue sections were incubated with citrate-based RNAscope 2.5 vs Target Retrieval buffer, pH 6.0 (ACD Bio; 322221), and then heated to 97 °C for 30 min before treatment with RNAscope 2.5 vs mRNA pretreat 3-Protease (ACD Bio; 322218) at 37 °C for 16 min. The human (RNAscope 2.5 VS Probe-Hs-MGLL; ACD Bio; 539159) and mouse (RNAscope 2.5 VS Probe-Mm-Mgll; ACD Bio; 478839) MAGL probes were then hybridized on the tissue for 60 min. To ensure tissue RNA integrity, a *UBC* positive control probe was used for human (RNAscope 2.5 VS Positive Control Probe-Hs-UBC; ACD Bio; 312029) and mouse (RNAscope 2.5 VS Positive Control Probe-Mm-Ubc; ACD Bio; 310779) specimens. A *dapB* negative control probe (RNAscope 2.5 VS Negative Control Probe_dapB; ACD Bio; 312039) corresponding to a bacterial gene not present in most mammalian tissue, was used for both species.

### Statistical analysis

GraphPad Prism v8.0 (GraphPad Software, La Jolla, CA, USA) was used for all statistical analysis. Statistical tests were used for each data set as indicated in the figure legends and all data are presented as means ± standard deviations (SD) with asterisks to indicate significant differences (ns = not significant, *p < 0.05, **p < 0.01, ***p < 0.001, ****p < 0.0001).

## RESULTS

### MAGLi 432 selectivity and potency in human and mouse brain lysates

MAGLi 432 [33] (**Fig. 1a**) is a non-covalent MAGL inhibitor that binds to the human (**Fig. 1b**) and mouse (not shown) MAGL enzymes. It binds with high affinity to the MAGL active site, with IC_50_ values of 4.2 nM for the human enzyme and 3.1 nM for the mouse enzyme as observed in enzymatic assays [33]. To investigate the selectivity and target occupancy of MAGLi 432 in mouse and human brain lysates, we carried out competitive activity based protein profiling (ABPP) with DMSO (control), 10 μM MAGLi 432 or 10 μM of the established covalent MAGL inhibitor JZL 184 (IC_50_ = 8.1 and 2.9 nM for human and mouse MAGL, respectively) (7,12). The lysates were then incubated with TAMRA-FP, a fluorophosphonate probe that binds irreversibly to serine residues in the MAGL active site, thus revealing whether MAGLi 432 selectively engages with MAGL by competitively inhibiting TAMRA-FP binding (**Fig. 1c,f**). Competitive ABPP was also carried out with a MAGL-specific probe [41] under the same conditions (DMSO, MAGLi 432 and JZL 184) to determine the specificity of MAGLi 432 for MAGL (**Fig. 1c-h**). In both experiments, we observed the almost complete absence of TAMRA-FP and MAGL probe fluorescent bands at ^~^33 and ^~^35 kDa in human (**Fig. 1c-e**) and mouse (**Fig. 1f-h**) lysates treated with MAGLi 432, suggesting that MAGLi 432 achieves an effective blockade of active MAGL. Significantly, MAGLi 432 selectively blocked binding of TAMRA-FP to MAGL but not to the active sites of other serine hydrolases. In contrast, JZL 184 only partially inhibited MAGL (83% and 78% decreases in human and mouse, respectively) compared to 100% inhibition for MAGLi 432, in all cases relative to the DMSO control. We also observed the non-specific binding of JZL 184 in mouse brain lysates revealed by non-specific signal inhibition at ^~^80 kDa (**Fig. 1f**). Normalized band intensities from both probes in human (**Fig. 1d,e**) and mouse (**Fig. 1g,h**) brain lysates were normalized to the total protein signal (**Fig. S1**). Overall, these results suggest that MAGLi 432 displays selectivity for MAGL over other serine hydrolases, in contrast to the irreversible inhibitor JZL 184.

**Figure 1.**
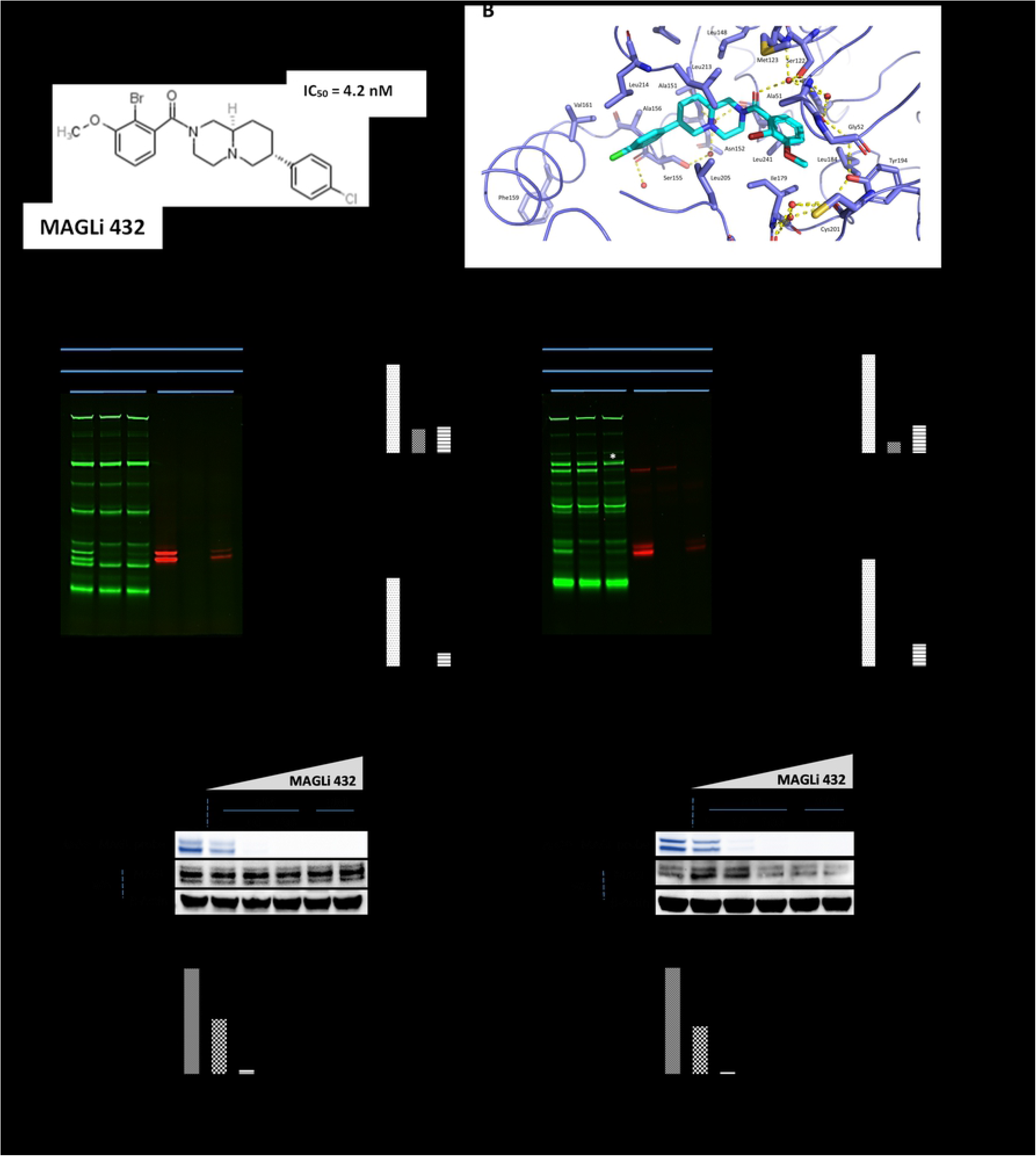
MAGLi 432 blocks MAGL in human and mouse brain lysates in a selective and potent manner. **(A)** Chemical structure of MAGLi 432 with an IC_50_ = 4.2 nM in human MAGL as determined by enzymatic assays. **(B)** X-ray co-crystal structure of MAGLi 432 (cyan) with human MAGL (blue). View of the active site of human MAGL (blue) reveals a non-covalent binding of the inhibitor. Interactions of MAGLi 432 with the catalytic Ser122 are mediated through the amide carbonyl group via a water molecule located in the oxyanion hole. Hydrogen bonds are depicted as yellow dashed lines. Selective MAGL inhibition by MAGLi 432 in human **(C-E)** and mouse **(F-H)** brain lysates was determined by competitive activity based protein profiling (ABPP). Brain lysates were incubated with DMSO, 10 μM MAGLi 432 or 10 μM JZL 184 and then with either a broad serine hydrolase activity-based probe (TAMRA-FP, green) or the MAGL-specific probe (red) before samples were loaded on a gel. Green fluorescent bands reveal serine hydrolases not inhibited by MAGLi 432. Red fluorescent bands reveal active MAGL not blocked by MAGLi 432. The normalized signal intensity was quantified as the average signal intensity for each MAGL fluorescent band (green or red) divided by the total protein signal intensity (Coomassie Brilliant Blue staining, **Fig. S1**) in human **(D,E)** and mouse **(G,H)** brain lysates (n = 2). **(I-L)** Assessment of MAGL potency was measured by competitive ABPP after incubation with ascending doses of MAGLi 432 in human **(I)** and mouse brain lysates **(J)** followed by incubation with the MAGL-specific probe. **(K, L)** The average signal intensities of active MAGL and total MAGL protein in human **(K)** and mouse **(L)** brain lysates were quantified as the total detectable active MAGL signal (ABPP) divided by the total the MAGL protein/β-actin signal (n = 2).

We determined the potency of MAGLi 432 by competitive ABPP in human and mouse brain lysates treated with ascending doses of the inhibitor (10 nM, 100 nM, 1 μM and 10μM), followed by incubation with the MAGL-specific ABPP probe. MAGLi 432 engaged MAGL in a dose-dependent, equipotent manner (**Fig. S2**), with an IC_50_ of 1–10 nM and complete inhibition at 100 nM in human (**Fig 1i,k**) and mouse (**Fig 1j,l**) brain lysates. These data confirm that MAGLi 432 is a highly selective, potent MAGL inhibitor (superior to JZL 184) that can occupy and engage with the human and mouse enzymes.

### MAGL is expressed and active in human and mouse brain tissue and NVU cells

Next we investigated the expression and activity of MAGL in the cortical regions of the brain, which feature the highest density of brain capillaries and NVU (43). *MAGL* mRNA expression was confirmed by *in situ* hybridization using a *MAGL*-specific probe on human and mouse cortical tissue slices (**Fig. 2a**). *MAGL* mRNA was broadly distributed across cortical tissue cells and its perinuclear arrangement suggested a cytoplasmic localization. In terms of cell-specific expression in the neurovasculature, *MAGL* mRNA was detected in close proximity to, but not directly co-localized with CD31^+^ endothelial cells, suggesting that MAGL may be expressed in closely associated mural cells, astrocytes and pericytes, which envelop BMECs within the vascular barrier.

**Figure 2.**
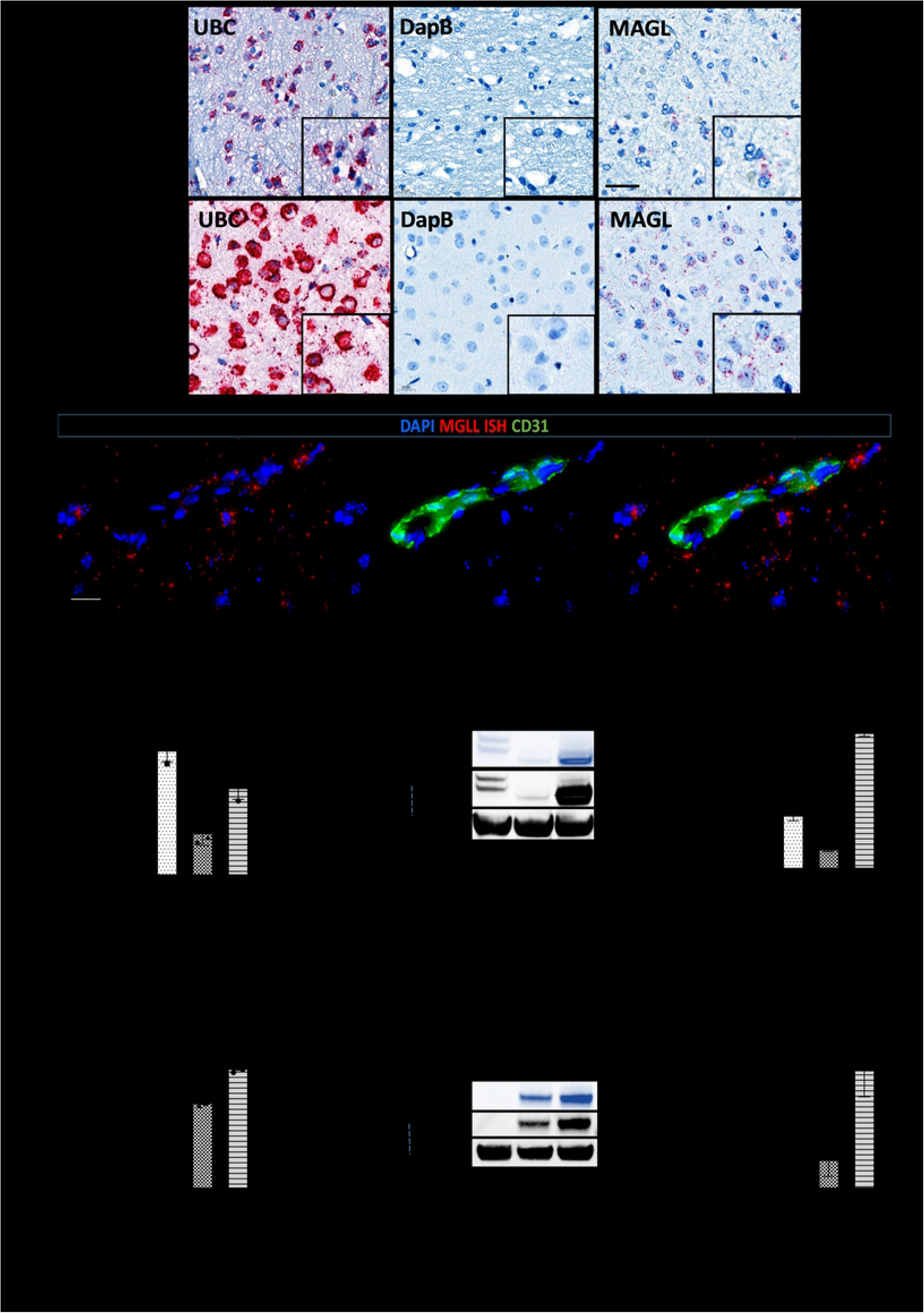
MAGL is expressed in the human and mouse cortex and is active at the level of the NVU (endothelial cells, astrocytes and pericytes). **(A)** Confirmation of *MAGL* mRNA expression by *in situ* hybridization (RNAScope) in human and mouse cortical brain tissue sections. Representative images show human and mouse brain cortical sections hybridized with *MAGL* RNAScope probes, then treated with chromogenic Fast Red substrate used for the detection of *UBC* (positive control for RNA integrity), *dapB* (negative control) and *MAGL* mRNA. Tissue sections counterstained with hematoxylin (n = 2). Scale bar = 20 μm. *MAGL* mRNA is expressed in close proximity to but does not co-localize with endothelial cells. **(B)** Immunofluorescence of mouse brain tissue sections (previously hybridized with *MAGL* mRNA) stained with an anti-CD31 antibody (green) to label endothelial cells (n = 2). Scale bar = 20 μm. MAGL is expressed and active in NVU cells. Human **(C-E)** and mouse **(F-H)** BMECs (hCMEC/D3 or pMBMEC), primary astrocytes (pHA or pMA) and primary pericytes (pHP or pMP) were used to assess *MAGL* expression within the NVU. Relative *MAGL* gene expression levels in human **(C)** and mouse **(F)** NVU cells measured by droplet digital qRT-PCR. Relative *MAGL* gene expression is normalized to housekeeping gene *EMC7* (n = 3, mean ± SD, one-way ANOVA with Tukey’s *post hoc* test, ns = not significant, *p < 0.05, **p < 0.01, ***p < 0.001, ****p < 0.0001). MAGL activity (ABPP) and total MAGL protein (western blot) in human **(D, E)** and mouse **(G, H)** NVU cells. Active MAGL was quantified as the average intensity of the total detectable MAGL signal divided by total MAGL protein/β-actin signal (n = 2, mean ± SD, one-way ANOVA with Tukey’s *post hoc* test, ns = not significant, *p < 0.05, **p < 0.01, ***p < 0.001, ****p < 0.0001).

To confirm these findings, we profiled MAGL expression in human and mouse NVU cells *in vitro*, including human primary astrocytes and pericytes and hCMEC/D3 BMECs, as well as mouse pMBMECs, primary astrocytes and primary pericytes. *MAGL* gene expression was assessed by droplet digital qRT-PCR and MAGL protein expression was assessed by ABPP and western blot (**Fig. 2c-h**). *MAGL* gene expression correlated with MAGL protein expression in each cell type. The analysis of human NVU cells (**Fig. 2c**) revealed that primary pericytes expressed the highest levels of MAGL, followed by BMECs and primary astrocytes (**Fig. 2c-e, Fig. S3a,c**). In contrast, whereas primary pericytes also exhibited the highest *MAGL* expression levels among mouse NVU cells, astrocytes were next, with pMBMECs accumulating little to no detectable MAGL (**Fig. 2f-h, Fig. S3b, d**). We concluded that *MAGL* is expressed in the mouse and human brain cortex and at the level of the NVU cells, and showed for the first time that pericytes express the highest levels of *MAGL* mRNA and MAGL protein in the NVU.

### MAGLi 432 triggers the robust elevation of 2-AG levels in human NVU cells

Having demonstrated an efficient blockade of MAGL activity in brain lysates using MAGLi 432, we next investigated the efficacy of MAGLi 432 *in vitro* by dose-dependent competitive ABPP on human BMECs (hCMEC/D3) as well as primary human astrocytes and pericytes. Cells were treated with increasing doses of MAGLi 432 (10 nM, 100 nM, 1 μM and 10 μM) or DMSO (solvent control) for 6 h. ABPP showed effective MAGL target occupancy (**Fig. 3a-b, Figs S4 and S5**) with *in vitro* IC_50_ values < 10 nM. Unsurprisingly, a slightly higher concentration of MAGLi 432 was required to fully block active MAGL in pericytes given these showed the highest level of expression among NVU cells. These results confirmed the *in vitro* potency of MAGLi 432 and revealed that 1 μM is an effective *in vitro* concentration for MAGL inhibition studies.

**Figure 3.**
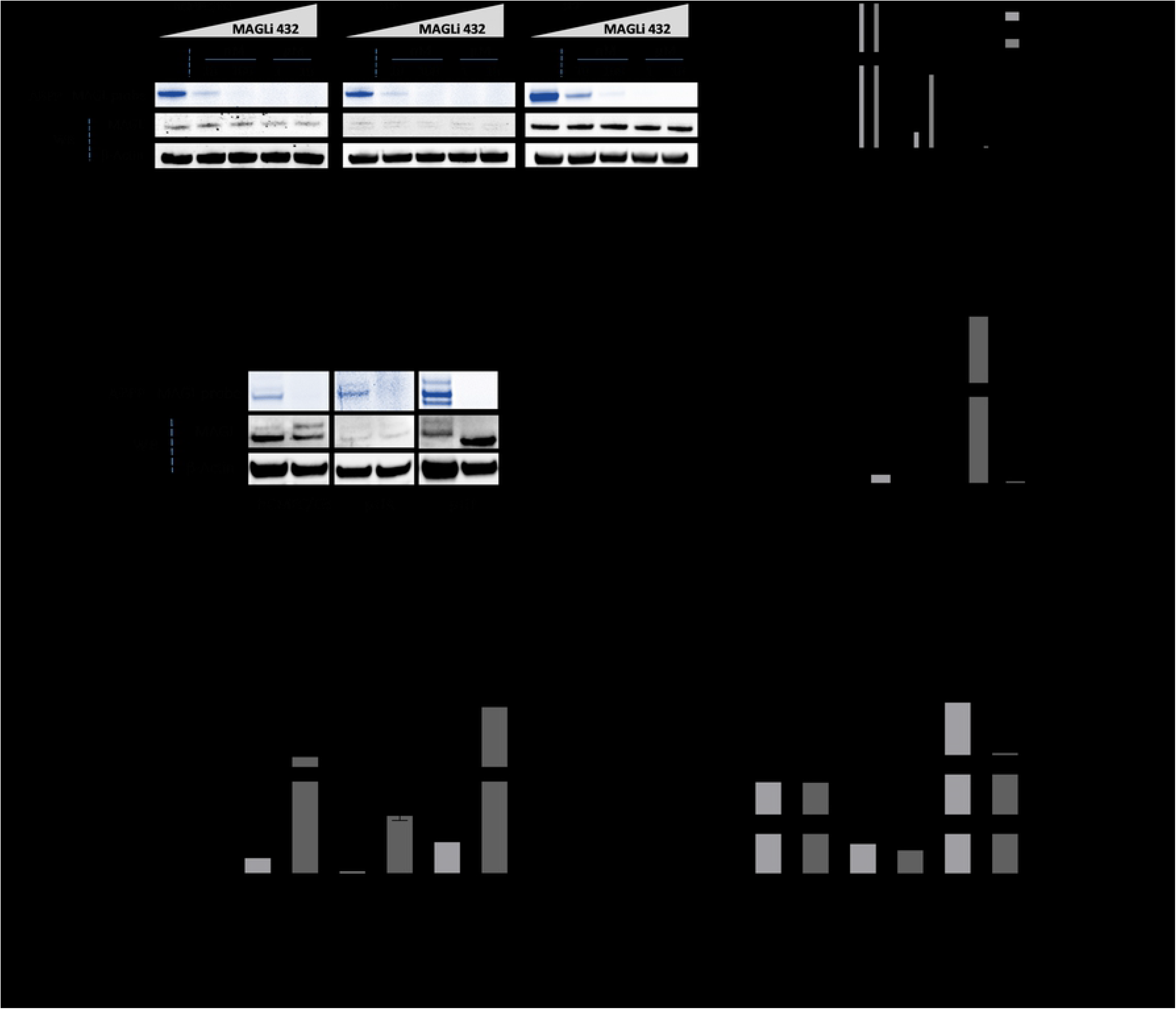
MAGLi 432 inhibits MAGL activity and robustly enhances 2-AG levels in human NVU cells. **(A)** Dose-dependent inhibition of active MAGL by MAGLi 432 in human BMECs (hCMEC/D3), astrocytes (pHA) and pericytes (pHP) *in vitro* assessed by ABPP and western blot applied to cell lysates 6 h after exposure to ascending doses of MAGLi 432 (10 nM, 100 nM, 1 μM and 10 μM) or DMSO. **(B)** The average signal intensities of MAGL activity and total protein were quantified as the total detectable active MAGL signal (ABPP) divided by the total MAGL/β-actin signal, with the highest signal normalized to 100%. **(C-D)** Assessment of a functional blockade of MAGL by MAGLi 432 in BMECs, astrocytes and pericytes *in vitro* by ABPP and western blot applied to cell lysates 6 h after exposure to DMSO or 1 μM MAGLi 432. The average signal intensities of MAGL activity and total protein were quantified as the total detectable active MAGL signal (ABPP) divided by the total MAGL/β-actin signal. **(E,F)** LC-MS measurement of 2-AG **(C)** and arachidonic acid (AA) **(D)** levels in cell lysates after treatment for 6 h (DMSO or 1μM MAGLi 432). Data are means ± SD, one-way ANOVA (ns = not significant, *p < 0.05, **p < 0.01, ***p < 0.001, ****p < 0.0001).

Given these findings, we investigated whether MAGLi 432 modulates the 2-AG/arachidonic acid axis *in vitro* by treating hMCEC/D3 cells as well as primary human astrocytes and pericytes with 1 μM MAGLi 432 or DMSO (solvent control) for 6 h. Again, target occupancy was measured by ABPP/western blot, and target engagement (via 2-AG/arachidonic acid modulation) was measured by LC-MS. ABPP showed that MAGLi 432 effectively blocked active MAGL, reducing it to undetectable levels in all cell types compared to the DMSO control (**Fig. 3c-d**). Target engagement studies revealed that MAGLi 432 increased 2-AG levels in all cell types (**Fig. 3e**), with the greatest effect (^~^70-fold increase) in pericytes but also an ^~^18-fold increase in BMECs and astrocytes. Interestingly, MAGLi 432 modulated arachidonic acid levels in a cell specific-manner, with no effect in BMECs, but significant depletion in astrocytes and pericytes (**Fig. 3f**). These data show that MAGLi 432 can modulate 2-AG levels in all cells, whereas the effect on arachidonic acid is cell-specific, suggesting that 2-AG and MAGL activity do not regulate arachidonic acid production *in vitro* in BMECs.

Next, we investigated whether inflammatory stimuli affect MAGL expression and activity in NVU cells. LPS administration leads to a significant increase in brain PGE_2_ levels *in vivo* [42], so we considered whether LPS could change the 2-AG/arachidonic acid balance in NVU cells in a MAGL-dependent manner. We treated BMECs, astrocytes and pericytes with LPS or HBSS (the LPS solvent) for 6 h, then measured MAGL expression by ABPP/western blot (**Fig. 4a,b**) and the modulation of 2-AG/arachidonic acid by LC-MS (**Fig. 4c,d**). The stimulation of BMECs with LPS did not alter MAGL expression. However, 50 and 100 ng LPS reduced the levels of active (**Fig. 4b**) and total MAGL (**Fig. S6b**) in astrocytes. Conversely, LPS increased the levels of both active (**Fig. 4b, Fig. S6a**) and total (**Fig. S6b**) MAGL in pericytes. Despite the changes in MAGL expression and activity in astrocytes and pericytes, LPS did not alter the cellular levels of 2-AG. However, arachidonic acid levels increased in response to 100 ng LPS in hCMEC/D3 cells and pericytes, and in response to 50 ng LPS in hCMEC/D3 cells, suggesting that 2-AG may not be the main source of arachidonic acid in those cells during inflammation. LPS significantly reduced arachidonic acid levels in astrocytes compared to the HBSS control, showing that LPS only induces the depletion of arachidonic acid in astrocytes.

**Figure 4.**
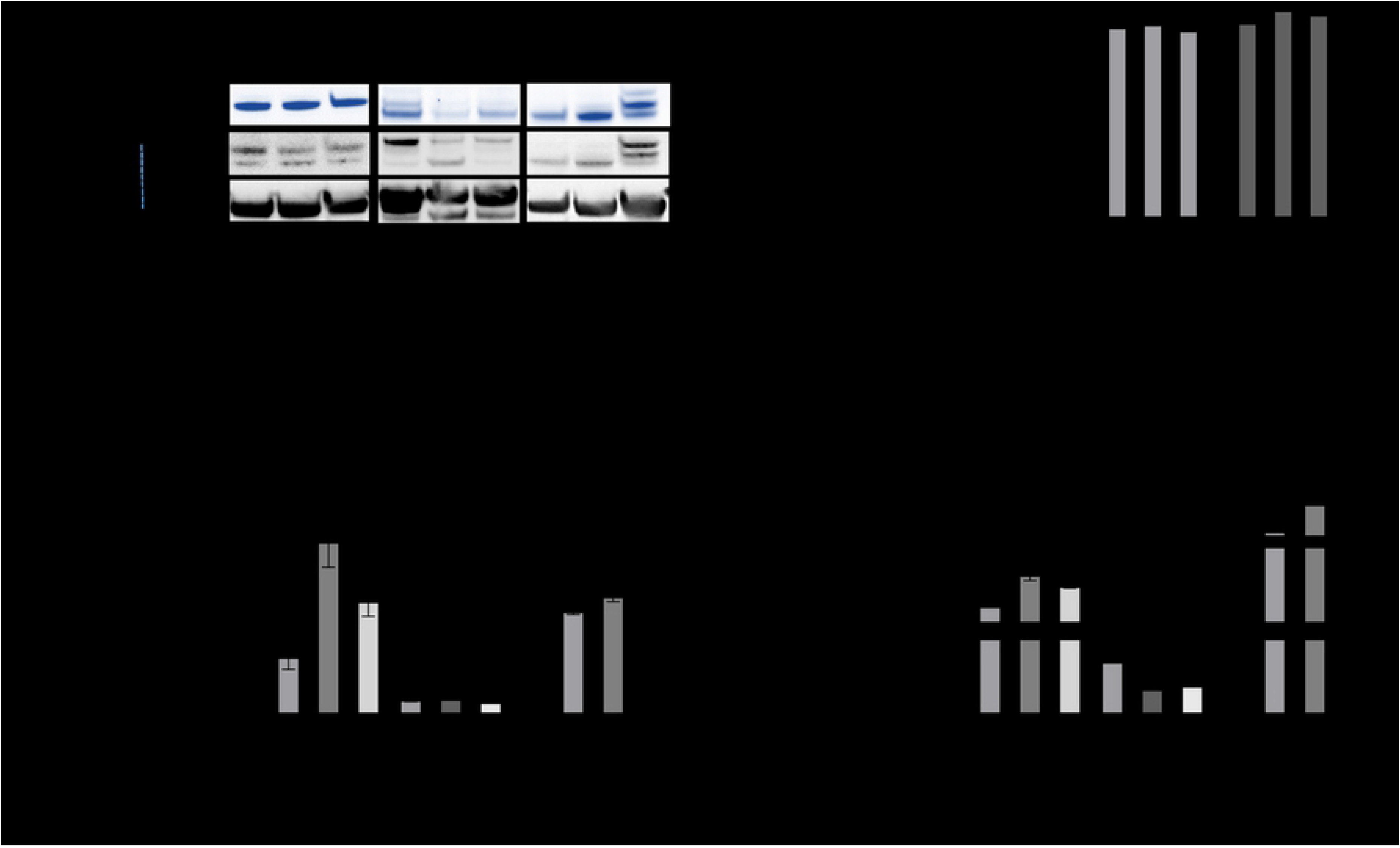
LPS does not modulate 2-AG in human NVU cells, but differentially regulates arachidonic acid levels in a cell-specific manner. **(A-B)** *In vitro* effects of LPS on active and total MAGL expression in BMECs as well as primary human astrocytes (pHA) and pericytes (pHP). (**A**) NVU cells incubated with HBSS (solvent) or 50 or 100 ng LPS for 6 h followed by ABPP and western blot applied to the cell lysates. (**B**) The average signal intensities of MAGL activity and total protein were quantified as the total detectable active MAGL signal (ABPP) divided by the total MAGL/β-actin signal. (**C, D**) LC-MS analysis of 2-AG **(C)** and arachidonic acid **(D)** levels in cell lysates after treatment for 6 h (HBSS or 50 or 100 ng LPS). Data are means ± SD, one-way ANOVA (ns = not significant, *p < 0.05, **p < 0.01, ***p < 0.001, ****p < 0.0001).

### MAGLi 432 achieves target occupancy and target engagement in a mouse model of LPS-induced neuroinflammation

Inhibiting 2-AG hydrolysis in the brain after injury or inflammation can limit the accumulation of arachidonic acid and thus restrict prostaglandin production. To investigate the ability of MAGLi 432 to achieve this effect *in vivo*, male CD-1 mice were randomized into three treatment groups (NaCl + vehicle, LPS + MAGLi 432 and LPS + vehicle) and were treated on 3 consecutive days with either NaCl or 1 mg/kg LPS, followed by either a vehicle solution or 1 mg/kg MAGLi 432 after a further 30 min (**Fig. 5a**). The assessment of cortical brain homogenates by ABPP and western blot revealed that a 1 mg/kg dosing regimen is sufficient to target MAGL activity, reducing it to almost undetectable levels (**Fig. 5b-c**). We also found that blocking active MAGL does not alter total MAGL protein expression in the brain and that LPS-induced inflammation does not modify active or total MAGL protein expression under our experimental conditions (**Fig. 3b-c**).

**Figure 5.**
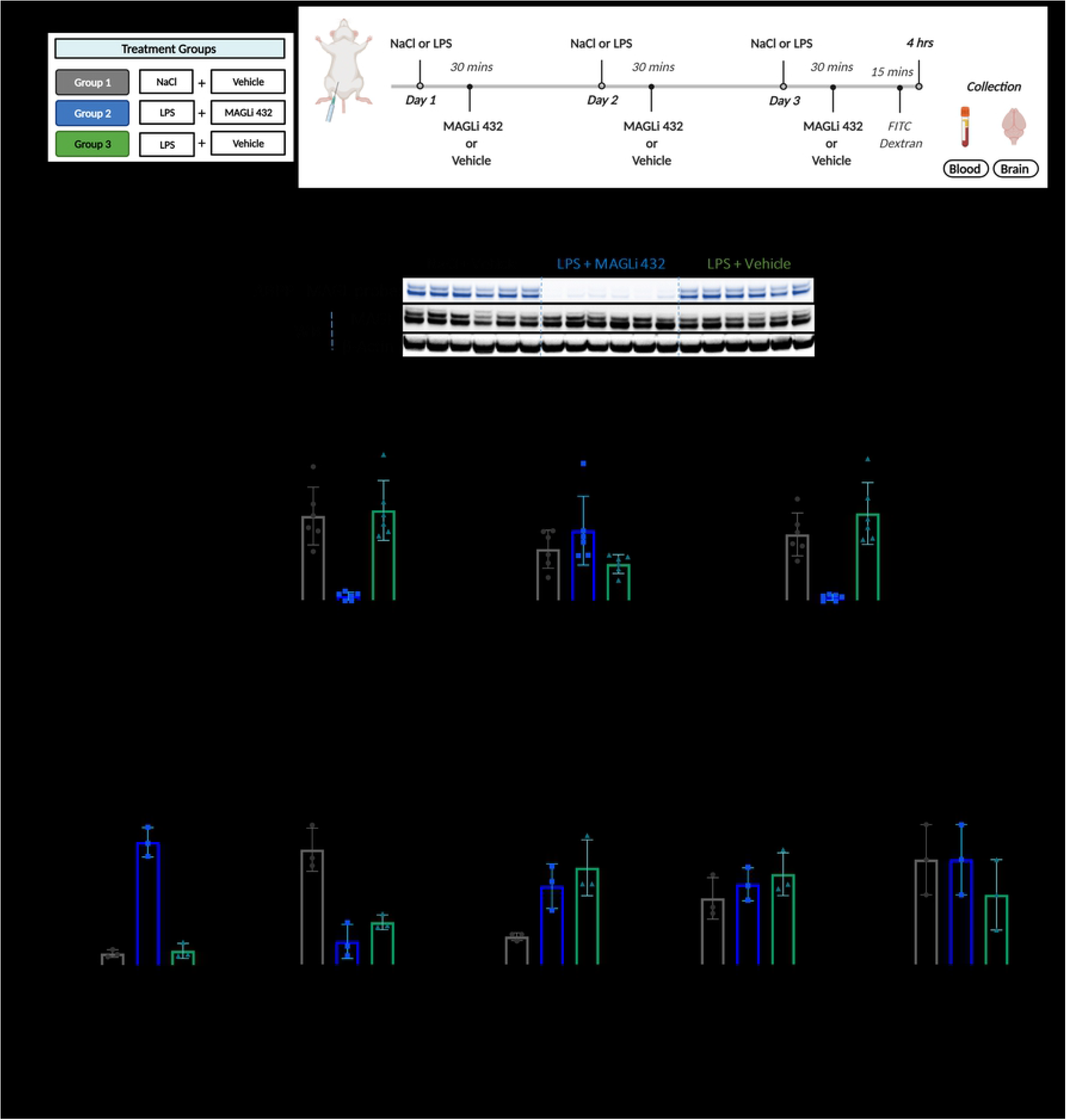
Target occupancy and engagement assays show that MAGLi 432 potently reduces MAGL activity in a mouse model of LPS-induced neuroinflammation. **(A)** Treatment groups and injection schedule of LPS and MAGLi 432. CD-1 mice were challenged with LPS or NaCl followed by MAGLi 432 or vehicle (i.p. administration over 3 consecutive days). Blood and brain tissue were collected for analysis. **(B,C)** ABPP and western blot analysis of cortical brain lysates from each treatment group reveal that MAGLi 432 achieves an effective blockade in the brain (n = 6). **(C)** The average signal intensities of MAGL activity and total protein were quantified as the total detectable active MAGL signal (ABPP) divided by the total MAGL/β-actin signal. (**D**) LC-MS analysis of 2-AG, arachidonic acid, PGE_2_, PGD_2_ and AEA in cortical lysates of each treatment group (n = 3). Data are means ± SD, one-way ANOVA with Tukey’s *post hoc* test, ns = not significant, *p < 0.05, **p < 0.01, ***p < 0.001, ****p < 0.0001).

We then used LC-MS to assess target engagement on brain cortical tissue to determine the levels of 2-AG, arachidonic acid, PGE_2_, PGD_2_ and AEA (**Fig. 5d**). Mice receiving MAGLi 432 accumulated ^~^10-fold more 2-AG than vehicle controls (mean increase = ^~^70 pmol/mg 2-AG in the LPS + MAGLi 432 treated group vs. 7–8.5 pmol/mg in the NaCl + vehicle and LPS + vehicle groups). Therefore, LPS alone had no significant effects on 2-AG. Arachidonic acid levels were significantly lower in both LPS-treated groups (**Fig. 5d**) along with a concomitant increase in PGE_2_ levels. MAGLi 432 achieved the greatest reduction in LPS-induced PGE_2_ but not significantly compared to the LPS control group. Neither PDG_2_ nor AEA was modulated in any of the treatment groups, supporting the specificity of MAGLi 432 for MAGL over FAAH (**Fig. 5d**). Collectively, these data confirm that MAGLi 432 engages MAGL in the brain, suggesting that MAGL inhibition reduces arachidonic acid and PGE_2_ levels in our subchronic LPS paradigm.

MAGL inhibition and the depletion of arachidonic acid pools in the brain following inflammation or injury can reduce the abundance of pro-inflammatory cytokines such as IL-6 and IL-1β [13,43]. Endocannabinoid signaling has also been shown to favorably modulate BBB permeability brought on by systemic LPS-administration [13] and 2-AG accumulates after traumatic brain injury (10), so we investigated whether 2-AG accumulation and the proposed anti-inflammatory effects of MAGL inhibition prevent leakage through the BBB in our 3-day LPS model. On the third day of LPS administration, a 70-kDa FITC dextran tracer was administered intravenously 15 min before euthanization to trace vascular leakage into the brain parenchyma. Brain cortical tissue from mice in each group was collected and stained with a CD31-specific antibody to label the endothelial cells (**Fig. 6a**). Examination of the tissue revealed intravascular confinement of the dextran tracer in the untreated mice (vascular co-localization) but the extravasation of dextran in both LPS-treated groups (**Fig. 6a**). MAGLi 432 therefore does not prevent LPS-induced leakage of the 70-kDa tracers into the brain parenchyma.

**Figure 6.**
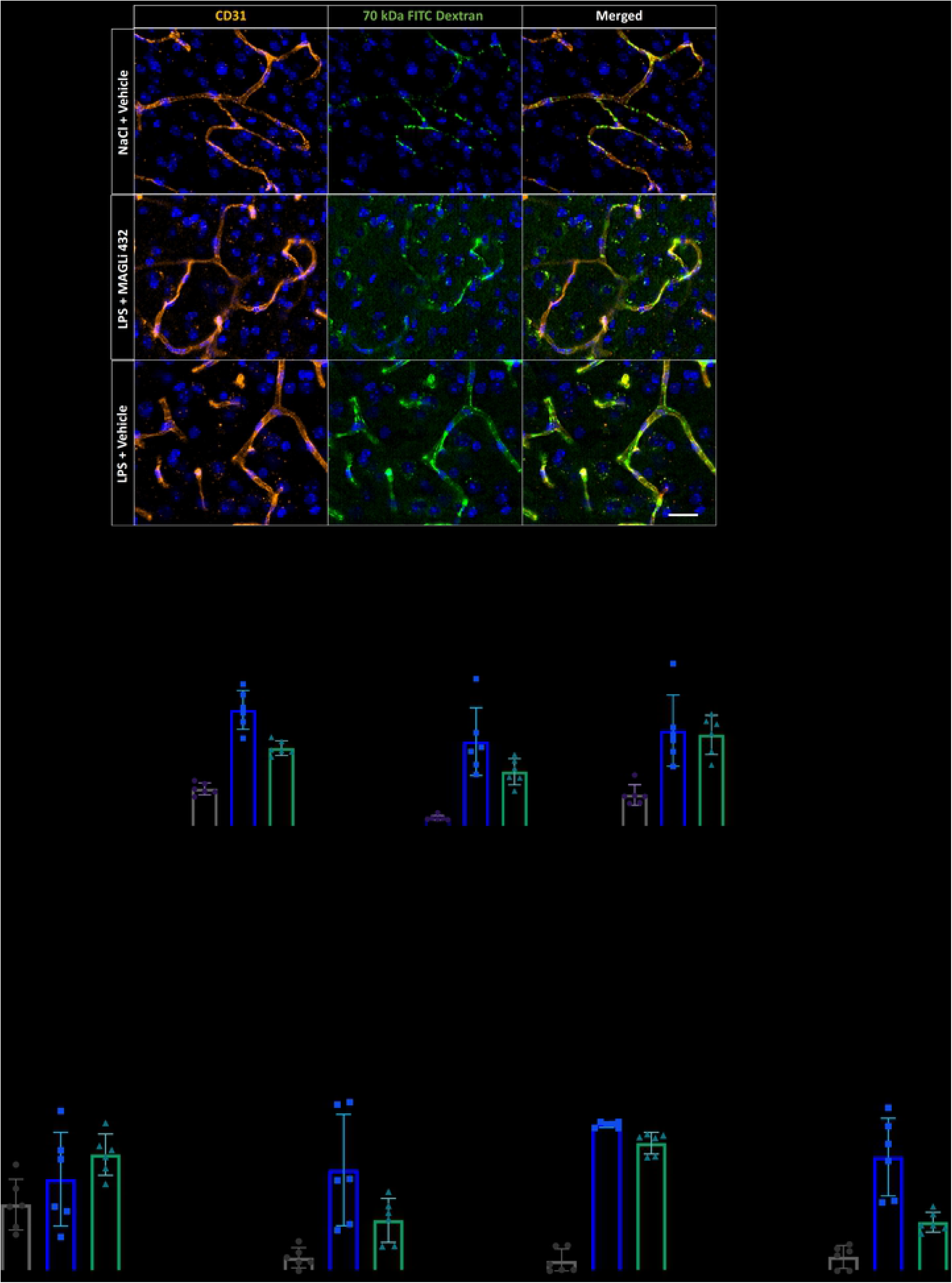
MAGLi 432 treatment after LPS challenge does not reduce BBB permeability and inflammatory cytokine expression in the cortex. **(A)** Immunofluorescence micrographs of 100-μm cortical sections from mice (n = 6) injected with 70-kDa FITC-dextran following LPS challenge and treatment (**Fig. 3A**) stained with an anti-CD31 (PECAM-1) antibody (orange) to label blood vessels and DAPI as a nuclear stain (blue). The experiment revealed no decrease in vascular permeability after exposure to MAGLi 432 compared to the LPS-only group (scale bar = 20 μm). **(B)** Assessment of vascular permeability by measuring fibrinogen extravasation in plasma and cortical brain lysates (n = 6, normalization to total protein determined with a BCA assay). **(C)** Relative expression of *IL-6, IL-1β, LCN2* and *TNF* genes assessed by droplet digital qRT-PCR. Gene expression was normalized to absolute gene expression levels per sample. Values represent results from two separate experiments. One-way ANOVA was used to compare the mean of each treatment group with the mean of every other group (n = 6, ns = not significant, *p < 0.1, **p < 0.01, ***p < 0.001, ****p < 0.00001).

We also measured the permeability of the BBB to fibrinogen, a 340-kDa blood-borne acute phase protein, by detecting it in the brain parenchyma. Fibrinogen is strongly upregulated in the circulation during inflammation and exacerbates inflammatory cascades following its extravasation across a disrupted BBB (44). We measured the ratio of cortical brain fibrinogen versus plasma fibrinogen, revealing a significant increase in the brain:plasma fibrinogen ratio after 3 days of LPS treatment, which was not ameliorated by MAGLi 432 (**Fig. 6b**). This confirms that MAGL inhibition does not prevent LPS-induced BBB permeability under our experimental conditions.

Finally, we determined the effects of our multi-dose LPS challenge on the inflammatory cytokine response in cortical brain lysates by droplet digital qRT-PCR. MAGLi 432 did not reduce the production of the pro-inflammatory cytokines IL-1β and IL-6 in response to LPS (**Fig. 6c**). Conversely, MAGLi 432 significantly increased LCN2 and TNF expression compared to the LPS treatment (**Fig. 6c**).

Taken together, our results suggest that MAGLi 432 neither prevented LPS-induced BBB permeability nor modulated the LPS-induced pro-inflammatory cytokine response in the LPS paradigm we used, despite robust MAGL target engagement and the accumulation of more 2-AG in the brain.

## DISCUSSION

In this study, we have shown that MAGLi 432 is a reversible, highly potent and selective MAGL inhibitor that can penetrate the brain *in vivo*. MAGL inhibitors have been developed for the therapeutic management of pain, movement disorders, and neurodegenerative diseases, involving mechanisms such as stimulating 2-AG production or limiting the expression of PGE_2_ and pro-inflammatory cytokines [11,13,44–46]. We focused on the characterization of MAGLi 432 and its ability to inhibit MAGL *in vivo*, its effect on LPS-induced BBB permeability, and its impact on NVU cells *in vitro*.

We found that MAGLi 432 is active against both human and mouse MAGL. The IC_50_ against human MAGL was 4.2 nM, comparable to that of irreversible inhibitors such as JZL 184 (IC_50_ = 8 nM), KLM29 (IC_50_ = 2.5 nM) and MJN110 (IC_50_ = 2.1 nM) [26]. The dose-dependent inhibition of MAGL activity without effects on MAGL protein levels was observed *in vitro* in pericytes, BMECs, and astrocytes, *ex vivo* in brain tissue homogenates, and *in vivo* in mouse brains. Only a few reversible MAGL inhibitors have been reported thus far, and MAGLi 432 may therefore have profound clinical implications. Specifically, the administration of irreversible inhibitors is associated with prolonged 2-AG elevation and the desensitization of CB receptors [29,47]. In contrast, reversible inhibitors should help to identify a therapeutic window of 2-AG elevation that elicits anti-inflammatory effects without CB receptor tolerance.

The high selectivity of MAGLi 432 for the active site of MAGL may also improve clinical outcomes by conferring a favorable safety profile. The selectivity of MAGLi 432 is not matched by the irreversible inhibitor JZL 184, as confirmed in our competitive ABPP experiments with the TAMRA-FP probe and a MAGL-specific probe. Furthermore, MAGLi 432 did not modify AEA levels in mouse brains *in vivo*, suggesting it does not inhibit FAAH (Ref). Irreversible MAGL inhibitors often lack selectivity, partially inhibiting FAAH, ABDH6 and ABDH12 and thus leading to off-target effects that hamper clinical development [27,48,49]. The importance of high selectivity was highlighted in a recent clinical phase I study of the FAAH inhibitor BIA 10-2474, which led to neurotoxicity that was fatal in one participant. ABPP selectivity assessment in human colon carcinoma cells revealed the binding of several additional serine hydrolases, which may have contributed to the reported adverse events [32,50].

The investigation of cell-specific MAGL expression and pharmacological modulation have often focused on CNS neurons and glial cells, with the neurovasculature receiving little attention [12,23,27,30,51–53]. We showed for the first time that MAGL is expressed in the microvasculature and in cells of the NVU. Cultured human and mouse brain microvascular endothelial cells, astrocytes and pericytes express MAGL, although pericytes are the predominant source of MAGL at the level of the neurovasculature. Accordingly, the inhibition of MAGL in pericytes may contribute most to therapeutic elevation of 2-AG levels in the brain, conferring neuroprotective functions including neurovascular integrity. Bulk RNA-Seq datasets often exclude mural cells from tissue extracts before sequencing, or their expression profiles are sequestered into broader cell groups, making it difficult to deduce their specific expression profiles. However, recent RNA-Seq datasets have begun to include these cells and their subsets, supporting our findings [54].

We then investigated the impact of MAGL inhibition on the levels of 2-AG and arachidonic acid, revealing a cell type-specific functional role. Specifically, we found that MAGL controls the 2-AG/arachidonic acid ratio in human peripheral astrocytes and pericytes, based on the observation that MAGLi 432 specifically raised 2-AG levels and slightly reduced arachidonic acid levels in these cells. However, MAGL inhibition in endothelial cells leads to a significant increase in 2-AG levels with no impact on arachidonic acid, suggesting that MAGL is not the main enzyme responsible for arachidonic acid production in these cells. In peripheral tissues, PLA_2_ is the main enzyme responsible for most arachidonic acid production and it may also play this role in brain microvascular endothelial cells. Indeed, the accumulation of arachidonic acid due to PLA_2_ activity in human brain endothelial cells and mouse models of vascular neuroinflammation has already been described [55,56].

The protective role of 2-AG helps to preserve BBB integrity following exposure to acute or chronic inflammatory insults [13,57]. We therefore sought to determine whether (i) inflammatory stimuli modulate MAGL expression and/or activity *in vivo* and (ii) MAGL inhibition could dampen LPS-induced BBB permeability and the production of pro-inflammatory cytokines. We observed no changes in active or total MAGL protein expression either *in vitro* or *in vivo* in response to LPS, indicating that MAGL expression is not modulated under inflammatory conditions in human NVU cells or in the mouse brain. Furthermore, multiple LPS doses over 3 days did not affect the abundance of total or active MAGL protein in the CNS. Although MAGL target engagement in the brain was significant, the inhibition of MAGL did not prevent LPS-induced BBB permeability. This was confirmed by the extravasation of high-molecular weight dextrans and fibrinogen into the brain parenchyma. Furthermore, the expression of key neuroinflammatory mediators, including IL-1β, IL-6, LCN2 and TNF, was either unaffected or even induced following the inhibition of MAGL.

Taken together, our data show that MAGLi 432 lacks anti-inflammatory effects in our experimental setting, which contrasts with multiple earlier studies reporting the *in vivo* anti-inflammatory effects of MAGL inhibition following a brain insult [10,11,13,45]. However, most of these studies involved irreversible inhibitors that lacked selectivity in models of acute inflammation. The binding and inhibition of additional endocannabinoid hydrolases such as FAAH and/or ABHD6/12 could achieve the observed protective effects by limiting the accumulation of agents that trigger 2-AG hydrolysis or the formation of arachidonic acid pools. Conversely, repetitive LPS injections may have created a hyper-inflammatory environment that is not representative of physiological inflammatory states. LPS is known to boost COX2 expression in the brain [58–60]. Although COX2 converts arachidonic acid into downstream prostaglandins, it can also directly oxidize 2-AG into prostaglandin glycerol esters, which are rapidly hydrolyzed into PGE_2_ or act directly on P2Y6 purinergic receptors [61–64]. These receptors are strongly upregulated in the vascular endothelium in the presence of LPS and increase the expression of pro-inflammatory cytokines and cell adhesion molecules [65,66]. Although MAGL inhibition can deplete pools of arachidonic acid and thus limit prostaglandin production, the LPS-related increase of COX-2 is not directly modulated by MAGL inhibitors and therefore does not ameliorate COX2 inflammatory pathways. Furthermore, cytokine expression is dynamic at the BBB, therefore snapshot measurements of may limit the scope of inflammatory modulation and determine when and where MAGL inhibition can be most effective in the restoration of vascular integrity.

## CONCLUSION

We have shown that the reversible inhibitor MAGLi 432 displays high selectivity and potency towards mouse and human MAGL. We observed the differential expression of MAGL in NVU cells and highlighted its relevance for the cell-specific production of arachidonic acid, which should lead to further investigations focusing on the role of MAGL in neurovascular insults. In future studies, more clinically relevant disease models involving dysregulated brain microvasculature should be used to investigate the effects of MAGL inhibition. For example, mouse models recapitulating disease pathology of AD (ex: APPP/PS1 transgenic mice)[67] or of MS (experimental autoimmune encephalomyelitis (EAE) mice) also develop dysregulation of the BBB due to disease-related inflammation[68]. Given the known neuroprotective role of 2-AG, our results suggest that NVU cells can produce beneficial levels of 2-AG, leading to the accumulation of protective secretory factors even during inflammation. Therefore, future studies with MAGLi 432 should investigate neuroprotective effects and different therapeutic dosing frequencies in acute and chronic preclinical models of neuroinflammation and vascular permeability. Experiments involving neurovascular cells derived from human induced pluripotent stem cells as well as 3D perfusable neurovascular models could provide further insights into the effects of MAGL pharmacological intervention on cell-cell dynamics within the NVU.

## ACKNOWLEDGEMENTS

We would like to thank Marie-Therese Miss for assistance with in vivo experiments, Agnes Nilly and Heinz Gutweiller for ordering and preparation of compound solutions for in vivo injections. Additionally, we would like to thank Florian Mohr for his assistance with ABPP and WB studies.

---

**Supplementary Table 1.** Data collection and refinement statistics for human MAGL compound 432 complex

|  | human MAGL<br>compound 432<br>complex |
| --- | --- |
| **Data collection** |  |
| Space group | C222<sub>1</sub> |
| Cell dimensions |  |
| *a*, *b*, *c* (Å) | 89.96, 127.45, 63.03 |
| $\alpha$, $\beta$, $\gamma$ (°) | 90, 90, 90 |
| Resolution (Å) | 1.16 (1.26-1.16) |
| *R*<sub>sym</sub> | 0.057 (0.80) |
| *I* / $\sigma I$ | 11.27 (1.09) |
| CC(1/2) | 0.999 (0.584) |
| Completeness | 99.8 (99.5) |
| Redundancy | 6.39 (5.99) |
| **Refinement** |  |
| Resolution (Å) | 63.72 – 1.16 |
| No. reflections | 122098 |
| *R*<sub>work</sub> / *R*<sub>free</sub> | 17.72/18.71 |
| No. atoms |  |
| Protein | 2306 |
| Water | 377 |
| Ligand | 28 |
| *B*-factors |  |
| Protein | 18.12 |
| Water | 34.30 |
| Ligand | 19.25 |
| R.m.s. deviations |  |
| Bond lengths (Å) | 0.008 |
| Bond angles (°) | 0.950 |

\*Values in parentheses are for highest-resolution shell.

**Supplementary Table 2.**
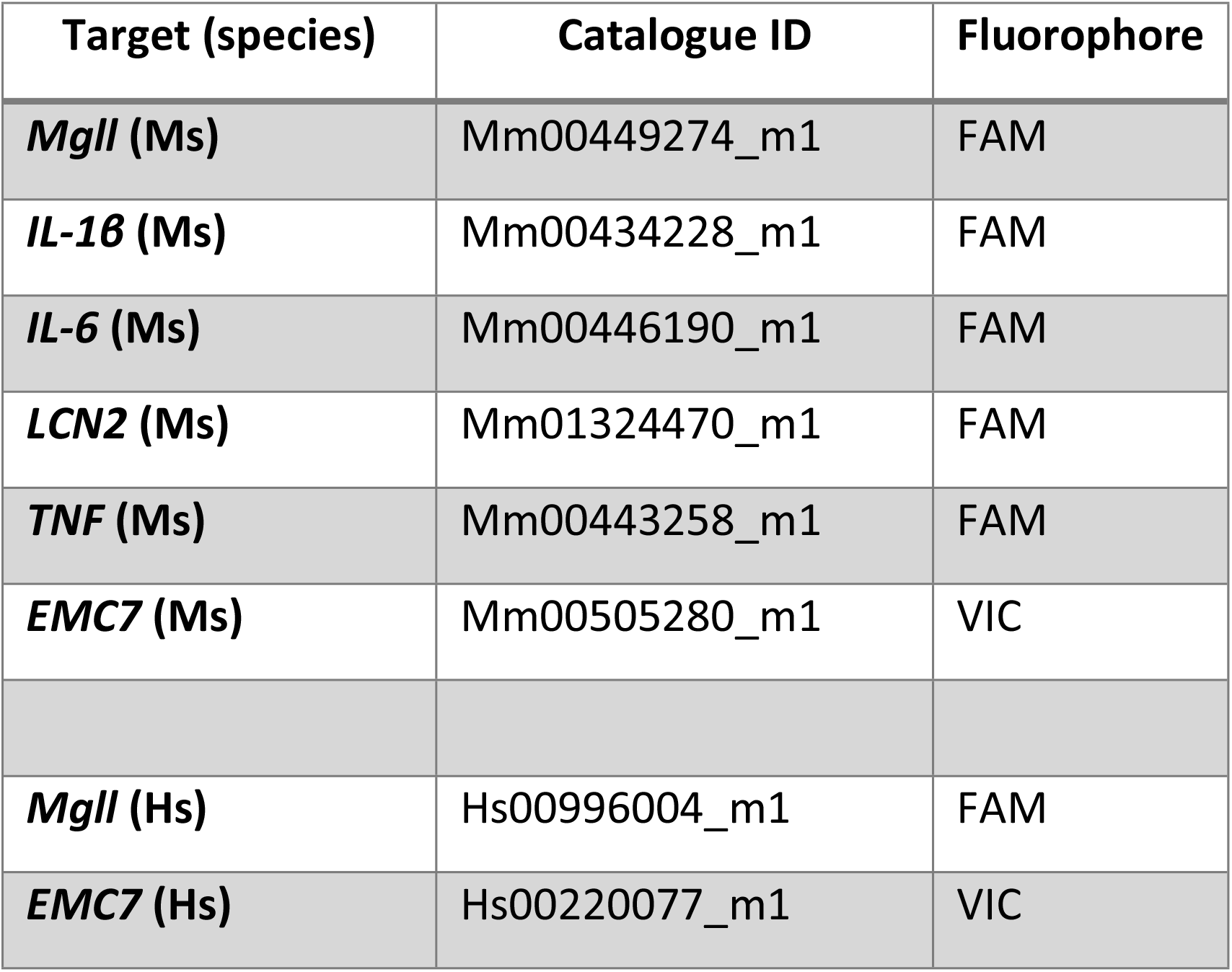
Droplet digital qRT-PCR TaqMan assays. Mm = *Mus musculus*, Hs = *Homo sapiens*.

**Figure S1. MAGLi 432 displays highly selective MAGL inhibition in human and mouse brain lysates.**

Selectivity of MAGLi 432 in human **(A)** and mouse **(B)** brain lysates was determined by gel based, competitive Activity Based Protein Profiling (ABPP). Brain lysates were incubated with either DMSO, 10 μM MAGLi 432 or 10 μM JZL 184 for 30 mins and then incubated with either broad serine hydrolase activity based probe, TAMRA-FP (green) or MAGL-specific probe (red) before samples were loaded on an SDS-PAGE gel and proteins separated by electrophoresis. Gels referenced from **Fig 1** were counterstained with Coomassie Brilliant Blue to visualize total protein per lane (n=2). Normalized MAGL activity quantified as the average signal intensity for each probe divided by total protein bands observed at the corresponding MW to MAGL bands (Coomassie Blue signal) (**Fig 1**).

**Figure S2. ABPP and WB quantitative analysis of MAGLi 432 dose-dependent potency in human and mouse brain lysates** (**Fig 1 I, J**). Assessment of MAGLi 432 potency was measured by incubation of ascending doses of MAGLi 432 in human brain lysates **(A, C)** and mouse brain lysates **(B,D)** as measured by competitive ABPP with the MAGL-specific probe. Average signal intensity of active MAGL and total MAGL protein in lysates quantified (**A,B**) as total detectable active MAGL band signal (ABPP) over total MAGL band signal (WB) or (**C,D**) total MAGL protein over total β-actin band signal (n =2).

**Figure S3.** Quantification of active MAGL (ABPP) and total MAGL protein (WB) in human (**Fig 2D**) and mouse (**Fig 2G**) NVU cells. Active MAGL was measured by incubation with the MAGL-specific probe via ABPP. Total MAGL protein expression and β-actin were determined by WB. Average signal intensity of active MAGL and total protein in lysates quantified (**A,B**) as total detectable active MAGL band signal (ABPP) over total MAGL band signal (WB) or (**C,D**) total MAGL protein over total β-actin band signal (n =2). Results are reported as mean ± SD, one-way ANOVA (ns = not significant, * = p < 0.05, ** = p < 0.01, *** = p < 0.001, **** = p < 0.0001).

**Figure S4.** Quantification of dose-response ABPP and WB to determine potency of MAGLi 432 in human NVU cells (BMECs, pHA and pHP) (**Fig 3 A, B**). Assessment of MAGLi 432 potency *in vitro* was measured by incubation of ascending doses of MAGLi 432 (10nM, 100nM, 1μM, 10μM) in human NVU Cells for 6 hours. Cell lysates from each group were then collected and then incubated with the MAGL-specific probe. Proteins were then separated by gel electrophoresis and in gel fluorescence was measured. Average signal intensity of active MAGL and total MAGL protein in lysates quantified (**A**) as total detectable active MAGL band signal (ABPP) over total MAGL band signal (WB) or (**B**) total MAGL protein over total β-actin band signal, with the highest signal normalized to 100%. (n =2).

**Figure S5.** Quantification of MAGL inhibition *in vitro* by MAGLi 432 in human NVU cells (hCMEC/D3, pHA and pHP) (from **Fig 3 C, D**). Assessment of MAGLi 432 *in vitro* was measured by incubation of DMSO (solvent control) or 1μM MAGLi 432 with human NVU cell culturesF for 6 hours. Cell lysates from each group were then collected and then incubated with the MAGL-specific probe. Proteins were then separated by gel electrophoresis and in gel fluorescence was measured. Average signal intensity of active MAGL and total MAGL protein in lysates quantified (**A**) as total detectable active MAGL band signal (ABPP) over total MAGL band signal (WB) or (**B**) total MAGL protein over total β-actin band signal. (n =2).

**Figure S6.** LPS modifies active and total MAGL protein expression in a cell-specific manner. Quantification of effects of HBSS (solvent control) and LPS (50ng and 100ng) *in vitro* by MAGLi 432 in human NVU cells (hCMEC/D3, pHA and pHP) (from **Fig 4 A, B**). Assessment of MAGLi 432 potency *in vitro* was measured by incubation of 1μM MAGLi 432 in human NVU Cells for 6 hours. Cell lysates from each group were then collected and then incubated with the MAGL-specific probe. Proteins were then separated by gel electrophoresis and in gel fluorescence was measured. Average signal intensity of active MAGL and total MAGL protein in lysates quantified (**A**) as total detectable active MAGL band signal (ABPP) over total MAGL band signal (WB) or (**B**) total MAGL protein over total β-actin band signal. (n =2).

